# The organotypic culture of mouse seminiferous tubules as a reliable methodology for the study of meiosis *in vitro*

**DOI:** 10.1101/2023.02.03.526942

**Authors:** P López-Jiménez, I Berenguer, I Pérez-Moreno, J.G. de Aledo, MT Parra, J Page, R Gómez

## Abstract

Male mouse meiosis has been traditionally studied using descriptive methods like histological sections, spreading or squashing techniques, which allow the observation of fixed meiocytes in either wildtype or genetically modified mice. For these studies, the sacrifice of the males and the extraction of the testicles are required to obtain the material of study. Other functional *in vivo* studies include the administration of intravenous or intraperitoneal drugs, or the exposure to mutagenic agents or generators of DNA damage, in order to study their impact on meiosis progression. However, in these studies, the exposure times or drug concentration are important limitations to consider when acknowledging animal welfare. Recently, several approaches have been proposed to offer alternative methodologies that allow the *in vitro* study of spermatocytes with a considerable reduction in the use of animals. Here we revisit and validate an optimal technique of organotypic culture of fragments of seminiferous tubules for meiotic studies. This technique is a trustable methodology to develop functional studies that preserve the histological configuration of the seminiferous tubule, aim homogeneity of the procedures (the use of the same animal for different study conditions), and allow procedures that would compromise the animal welfare. Therefore, this methodology is highly recommendable for the study of meiosis and spermatogenesis, while it supports the principle of 3R′s for animal research.

## 1. Introduction

Animal experimentation with mammals has been used in basic and applied research [1]. In particular, the house mouse, *Mus musculus*, is extensively used as mammalian model for experimental research, contributing to finding solutions to biological and biomedical questions related to human health. However, steaming from the proposal of Russell and Burch for the replacement, reduction and refinement of studies with life animals, the 3R’s rule [2], many laboratories, researchers and scientific agencies have encouraged in the past years the development of alternative methods. This is recommended not only by ethical reasons, avoiding suffering, pain, stress and/or death of animals, but also by the aim of achieving optimized *in vitro* methodologies that improve the quantity and quality of the research data.

The production of male gametes is called spermatogenesis, which is the transformation of spermatogonial diploid cells into haploid spermatozoa, is a process that occurs over an extended period of time inside the testis [3,4]. Spermatogenesis comprises three events: i) stem germ cells (spermatogonia) proliferation, ii) meiosis of spermatocytes, and iii) maturation of spermatids into spermatozoa (spermiogenesis) [5]. Meiosis itself implies two rounds of chromosome segregation with a single premeiotic event of DNA replication [6], giving rise to the production of haploid spermatids. These, later undergo dramatic morphological changes resulting into the maturation of fully competent motile spermatozoa [7].

At the histological level, spermatogenesis takes place in the seminiferous epithelium of the testis, a highly specialized cell environment. Growth factors and hormones tightly regulate many of these crucial steps leading to the formation of spermatozoa [5]. The organization of the seminiferous epithelium is unique since it is composed of a monolayer of the highly specialized Sertoli cells, which facilitate the progression of germ cells to spermatozoa via direct contact with them and by influencing both the development of spermatogonial germ cells and meiocytes [8,9]. Therefore, the histological conformation of the seminiferous epithelium is a key requirement of gamete production in mammals [5].

Male mouse meiosis has been traditionally studied using descriptive methods like cell spreading [10] or squashing [11], histological sections or electron microscopy. In virtually all cases, seminiferous tubules are freshly taken from sacrificed individuals and immediately processed. Nevertheless, several studies have pursued the development of *in vitro* systems. The first dated attempt was carried out in 1920 [12].

Later, many studies attempted to develop spermatogenesis *in vitro* by culturing dissociated testis cells. However, those studies reported that the entry of premeiotic cells into meiosis was blocked under *in vitro* conditions by erroneous or ineffective contact between Sertoli and germ cells [13]. Recent studies have progressed into this exciting context through the use of spermatogonial stem cells for *in vitro* spermatogenesis and potential *in vivo* applications for the restoration of fertility [14,15]. Moreover, key advances in biomedical research have permitted the application of *in vitro* mouse and human spermatogonial stem cells proliferation to achieve their transplantation into the seminiferous tubules of infertile mouse or men to restore fertility [16-18], yet the isolation of spermatogonial stem cells implies several complex procedures. For this reason, nowadays, the most successful results have been obtained from the organotypic culture of seminiferous tubules. In 2011, a hybrid gas-liquid interphase method was established by Sato and colleagues, where germ stem cells were capable of producing fertile spermatids and sperm *in vitro* [19]. In that publication, neonatal mouse testes containing primitive spermatogonia could produce spermatids and reproductively competent sperm *in vitro* with a serum-free culture media over an agarose gel. Those authors also combined the organ culture method with a spermatogonial stem cell transplantation technique [20,19]. This organotypic culture method set the basis for the use of *in vitro* controlled manipulation of the micro-environmental conditions required for proper murine spermatogenesis. Further studies succeeded in producing functional sperm from primitive spermatogonia in explanted neonatal mouse testis tissues [21]. Since then, several researchers have used this methodology to display functional approaches to the study of mouse spermatogenesis [22-30]. Latest advances using modified methodologies built upon the foundation of original protocols, have proposed that culturing isolated spermatogonia tubules, as opposed to tissue masses, offer distinct advantages in precisely evaluating the diverse environmental factors influencing the progression of spermatogenesis [31]. Alternatively, the polydimethylsiloxane-chip ceiling method is suggested to ensure a uniform supply of nutrients and oxygen throughout the testicular tissue [32]. Moreover, the integration of single polydimethylsiloxane-chip platforms, combined with real-time cellular paracrine support from allogeneic bone marrow mesenchymal stem cells, holds promise in providing a steady and stable microfluidic flow for the advancement of spermatogonial cultures [33]. Nonetheless, accomplishing a complete *in vitro* spermatogenesis capable of yielding a quantity of spermatozoa comparable to age-matched *in vivo* controls remains a significant technical challenge, especially in the context of human reproduction.

Besides the exciting advances for the biomedical applications of spermatogonia stem cells research as an approach to deal with human infertility, basic science also can benefit from the use of *in vitro* methodologies [34]. While the main aim of previous studies has been setting the conditions for successful production of spermatozoa, this can be also applied to more basic research, as for instance the study of meiosis. This special kind of cell division is crucial to increase the variability in the offspring. Furthermore, the proper occurrence of genetic and chromosomal events during meiosis is critical for the integrity of the genome that is transmitted from one generation to the next one. Indeed, many of the inherited chromosome aberrations are produced as defects in chromosome pairing, recombination or segregation during meiosis.

Contrary to other biological models in which meiotic cells manipulation in culture is feasible, study of meiosis in mammals has been partially obstructed by the technical difficulties to perform *in vitro* experiments. However, the ethical concerns in the application of physical or chemical treatment that can cause animal stress of pain, as well as to the large number of animals needed to carry out some experiments are pushing toward the development of alternative methods or the use of alternative models. Therefore, the implementation of *in vitro* techniques for the study of mammalian male meiosis is an interesting and necessary tool in meiosis research. In the past years we have successfully used some of these methods, being the organotypic culture of seminiferous tubules the most successful and reliable technique. We have particularly used the protocol described the Sato *et al*, with some specifications and modifications. Here we present a detailed description of the improved protocol that we have previously used [35-38], including specific tips and validation data, and also indicating some limitations of use.

## 2. Materials

### 2.1. Sample extraction

1. Testis from prepuberal or adult mice are extracted after euthanasia of the individuals, following local and regional ethics committees’ guidelines. Testis are detunicated with the help of precision tweezers in a glass petri dish with 1 ml of PBS 1x. Detunicated seminiferous tubules should remain in PBS 1x at 34 ºC until culture.
2. Seminiferous tubules are slightly disentangled with two forceps and cut transversally in small fragments with a sterile razorblade. These fragments are then be cultured in twelve-well plates and maintained at 34 ºC in an atmosphere with 5% CO for the duration of the study following the protocol explained below.

### 2.2. Agarose gel and media preparation

1. 20 ml of a 1.5% agarose gel are prepared by dissolving agarose in distilled water in a microwave. The agarose gel is then let to solidify in a 9 cm diameter glass petri dish.
2. Medium: MEMα culture medium supplemented with 10% KnockOut Serum Replacement (KRS) and 1% antibiotics (Penicillin/Streptomycin) is prepared and kept at 34 ºC until use.
3. For inhibition studies, the selected drug should be diluted in the culture medium at the correspondent concentration.

### 2.3. Slides preparations

1. Slides for the squashing technique: the slides are washed in a 1:1 ethanol-acetone solution and covered with poly-L-lysine (Sigma) to promote cell adhesion.
2. Slides preparation for the spreading technique: the slides are washed in a 1:1 ethanol-acetone solution and kept dry until use.

## 3. Methods

### 3.1. Organotypic cultures of control seminiferous tubules

1. 15x15mm sections of 3.2 mm thick agarose squares are placed in each well of the culture plate (Figure 1A) (see notes 1 and 2).
2. 2 ml of medium are placed in each well of a sterile twelve-wells culture plate (Figure 1A).
3. Culture plates are kept overnight at 4ºC to allow the interchange of water and MEMα between the liquid and agarose. After 24 hours, the medium of the wells is refreshened. For control studies, 600μl of fresh medium is added to each well. This should not cover the upper surface of the agarose gel.
4. The seminiferous tubules of freshly detunicated testis are cut in small pieces of approximately 5 mm length. The fragments of seminiferous tubules are placed over the half-soaked agarose gel and are kept in an air-liquid interphase (Figure 1B) (see note 3).
5. Well plates are kept at 34°C in an atmosphere with 5% CO for the duration of the study.
6. After the appropriate time, fragments of seminiferous tubules are moved to a Petri dish containing PBS 1x until used for squashing or spreading techniques.

**Figure 1:**
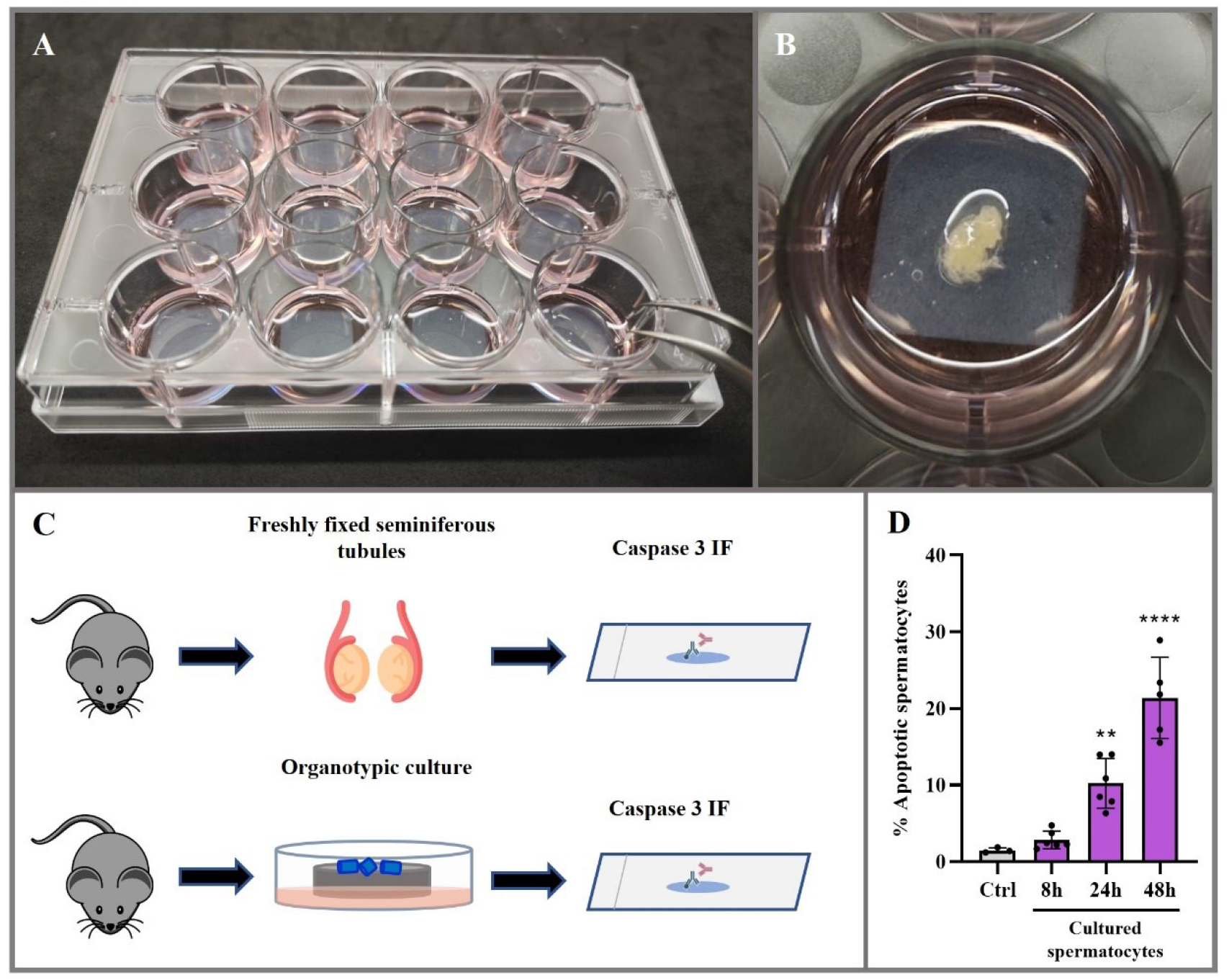
Organotypic cultures of control seminiferous tubules. (A) Photograph of a representative twelve-well culture plate containing medium and agarose gels. (B) Organotypic culture of seminiferous tubules in a selected well. The image shows the media, the agarose gel and the fragments of seminiferous tubules. (C) Experimental approach for the comparison of the level of apoptosis between freshly fixed seminiferous tubules extracted from dissected testis, and the cultured seminiferous tubules with fixation after culture. In both cases, immunolabelling of cleaved Caspase 3 was used to assess the extent of apoptosis. (D) Quantitative analysis of the percentage of apoptotic spermatocytes in freshly fixed seminiferous tubules and the cultured seminiferous tubules (at least three biological replicates for each condition, minimum of 500 spermatocytes per condition and individual). Data represents mean ± SD, p<0.05 (**), p. <0.005 (****), One-way ANOVA.

### 3.2. Inhibition studies with the organotypic culture method

1. To inhibit proteins implicated in meiosis progression in cultures of seminiferous tubules, the required concentration of the drug is usually between 10x to 100x higher that the concentration used for comparable studies with somatic cell cultures (see notes 5 and 6). This is due to the characteristics of the organotypic culture *per se*, as the drug diluted in the medium is absorbed through the agarose gel and across the seminiferous tubules before reaching the spermatocytes. This procedure is drastically different to adding drugs on top of somatic adherent cells cultures. Examples of these functional studies, where the inhibition of selected proteins *in vitro* in organotypic cultures is compared to the equivalent *in vivo* genetically modified model offers reliability to this method [35-38].
2. When designing the experimental approach, the optimized concentration of the drug needs to be identified after an exhaustive analysis of various concentrations. The duration of the treatment depends on the target spermatocytes to study. Previous reports about the duration of spermatogenesis and the overlapping waves of meiotic divisions should be taken into account [39,5].
3. When developing the organotypic culture for functional assays with the use of drugs or inhibitors, after 24 hours of media transfer to the agarose gel (see step 3.1/3), the media is replaced with the corresponding concentration drug of study diluted in 600μl of MEMα (see note 3).

Both control or treated spermatocytes of organotypic cultures can be processed by either the squashing or the spreading methodologies. The steps are provided below in sections 3.3. and 3.4.

### 3.3. Squashing of cultured seminiferous tubules

1. The cultured fragments of seminiferous tubules are moved to a new glass chamber containing a 2% PFA (BDH AnalaR) + 0,05%Triton X-100 (Sigma) fixation solution for 15 (see notes 1 and 2).
2. Preparation of squashed seminiferous tubules are performed following conventional previous protocols [11]. In brief, two or three 2cm length fragments of the fixed post-cultured seminiferous tubules are deposited in a slide that contains poly-L-lysine. The fragments of seminiferous tubules are carefully disintegrated with the aid of curved toothed forceps. Next, an additional drop of formaldehyde-Triton fixative was added, and a coverslip is placed on top of the material. Holding the coverslip from one of the corners, the fragments of the seminiferous tubules are softly squeezed by passing a pen over the coverslip. Then, the cells are squashed by applying pressure with the thumb on the coverslip to obtain a monolayer of cells.
3. The samples are then submerged in liquid nitrogen and stored at -80°C until future use. In case they are to be used immediately, the coverslip is removed with the help of a razor and slides are rehydrated in PBS. The rest of the preparations can be stored at -80°C for long periods of time. When needed, they should be immersed again in liquid nitrogen for a few seconds. Then, the coverslip is removed with the help of a razor and the slides are rehydrated in PBS. The following steps are equal to the fresh preparations produced the day of the organotypic culture procedure.
4. Immunolabelling methods of the squashed preparations can be done following standard protocols. In brief, the slides with the squashed seminiferous tubules are rinsed three times for 5 minutes in PBS and incubated 1 hour at 37ºC, or overnight at 4°C, with primary antibodies diluted in PBS. Then, the slides are rinsed three times for 5 minutes in PBS and incubated for 1 hour at room temperature with the secondary antibodies. Finally, the slides are counterstained with 10 μg/ml DAPI for three minutes, rinsed in PBS and mount with Vectashield (Vector Laboratories).

### 3.4. Spreading of spermatocytes extracted from cultures seminiferous tubules

1. Preparations of spreaded spermatocytes are performed following conventional previous protocols [10] (see notes 1 and 2). In summary, the post-cultured fragments seminiferous tubules are placed in a glass petri dish with 300 μL of PBS, minced with a scalpel, and disaggregated with two curved forceps. 3 mL of PBS are then added, and the solution is poured into a centrifuge tube. The content is resuspended with a Pasteur pipette and let to settle for 5 minutes to precipitate the larger fragments. The supernatant is then collected and transferred to a new tube that is centrifuged for 8 minutes at 1200 r.p.m. After that, most of the supernatant is discarded, leaving a small volume for resuspension with the rest of the material. The tube is then gently tapped to homogenize the mixture. Another 4 mL of PBS are added for a second centrifugation of 8 minutes at 1200 r.p.m. Supernatant is discarded, and 300 μL of 100 mM sucrose was added slowly to the pellet, resuspending every 3 drops. This cell suspension is then spread on slides that had been previously immersed in 1% paraformaldehyde in distilled water (pH=9) containing 0.15% of Triton X-100. 15μl of the suspension is added per slide and placed horizontally in a humid chamber. Preparations are kept in humid and dark conditions for 2 hours. Afterwards, the chamber cover is partially opened, and the samples were let to completely dry. After that, they are washed with 0.4% Photoflo (Kodak) in distilled water twice. Once dried, they are stored at -80°C, except those that are to be used immediately, which were rehydrated in PBS.
2. Immunolabelling methods of the squashed preparations can be done following standard protocols (explained in the previous section for the squashing technique).

### 3.5. Validation and reliability of the of organotypic culture method

*In vitro* studies are never fully equivalent to the *in vivo* condition as they lack histological and endocrine regulation, so that it is important to carry out validation tests. In order to verify if the organotypic culture method is trustworthy for meiotic studies, we have carried out three experiments that identify the level of suitability and reliability of this methodology.

1. First, we conducted a study to determine the level of spermatocyte apoptosis in organotypic cultured seminiferous tubules at different times. For this, we immunolabelled cleaved Caspase 3 and quantified spermatocytes with positive labelling in freshly fixed seminiferous tubules directly extracted from the testis (*in vivo* condition), and in squashed preparations of cultures spermatocytes at 8, 24 and 48 hours (*in vitro* condition) (Figure 1C). In all time points spermatocytes are still alive but increasing levels of apoptosis are observed with culture time (Figure 1D). Although the level of apoptosis after 48h is around 20% average, the spermatocytes that are not undergoing apoptosis show a normal pattern of distribution of the meiotic usual markers (discussed below). We conclude that the organotypic culture of seminiferous tubules is a trustable methodology for up to 48 hours. After that period, researchers should take into consideration potential side effects of the *in vitro* methodology.
2. Second, we conducted an experiment aiming to assess if organotypic cultured spermatocytes accumulate DNA damage as a consequence of the culture. To do this, we analyzed by immunolabelling the localization patterns of Synaptonemal Complex Protein 3 (SYCP3), as a marker of synapsis progression [40], and γH2AX, as a known marker of meiotic DNA double-strand breaks (DSBs) [41]. Results showed that DNA endogenous damage patterns are equivalent in spermatocytes directly extracted from the testis and in spermatocytes obtained from seminiferous tubules cultured for 24h (Figure 2A). We concluded that there is no significant difference between the pattern of distribution of the meiosis markers between the *in vivo* and the *in vitro* conditions, at least during the stages of prophase I and corroborated that culture do not induce new DSBs (Figure 2B).
3. A final experiment was conducted to address the suitability of the organotypic culture method for functional studies on DNA repair. To do this, we studied the effect of gamma radiation. We compared three different conditions (Figure 3A): a) 3 different mice were treated with 5 Gy of gamma radiation and sacrificed at 1, 6 or 24 hours after treatment, respectively; b) a mouse was irradiated similarly, immediately sacrificed and the seminiferous tubules maintained in culture and subsequently processed at 1, 6 or 24 hours; c) tubules of a control mouse were irradiated while in culture and subsequently processed at 1, 6 or 24 hours. The results showed that the three approaches were equally efficient to induce exogenous DNA damage, with no significant differences in the total number of induced γH2AX foci (Figure 3B, correspondent to condition c). Moreover, the dynamics of DNA repair is comparable in all the situations, with progressive reduction of γH2AX foci at 6 and 24 hours. Only a small difference could be observed 24 hours after irradiation, as both cultured conditions displayed a slightly higher number of remaining γH2AX foci (Figure 3C). Therefore, although the culture of the seminiferous tubules does not substantially perturb the dynamics of DNA repair after irradiation, the different physiological environment in this condition can introduce slight differences in the absolute number of DSBs recorded. This was expectable, since previous studies have reported a delayed progression of spermatogenesis in cultured seminiferous tubules [19,29].
4. In conclusion, the study of the level of apoptosis, synapsis progression, endogenous and exogenous DNA damage induction and repair dynamics allow us to conclude that the organotypic culture of seminiferous tubules is a trustable method to study male mouse meiosis *in vitro* (see note 7).

**Figure 2:**
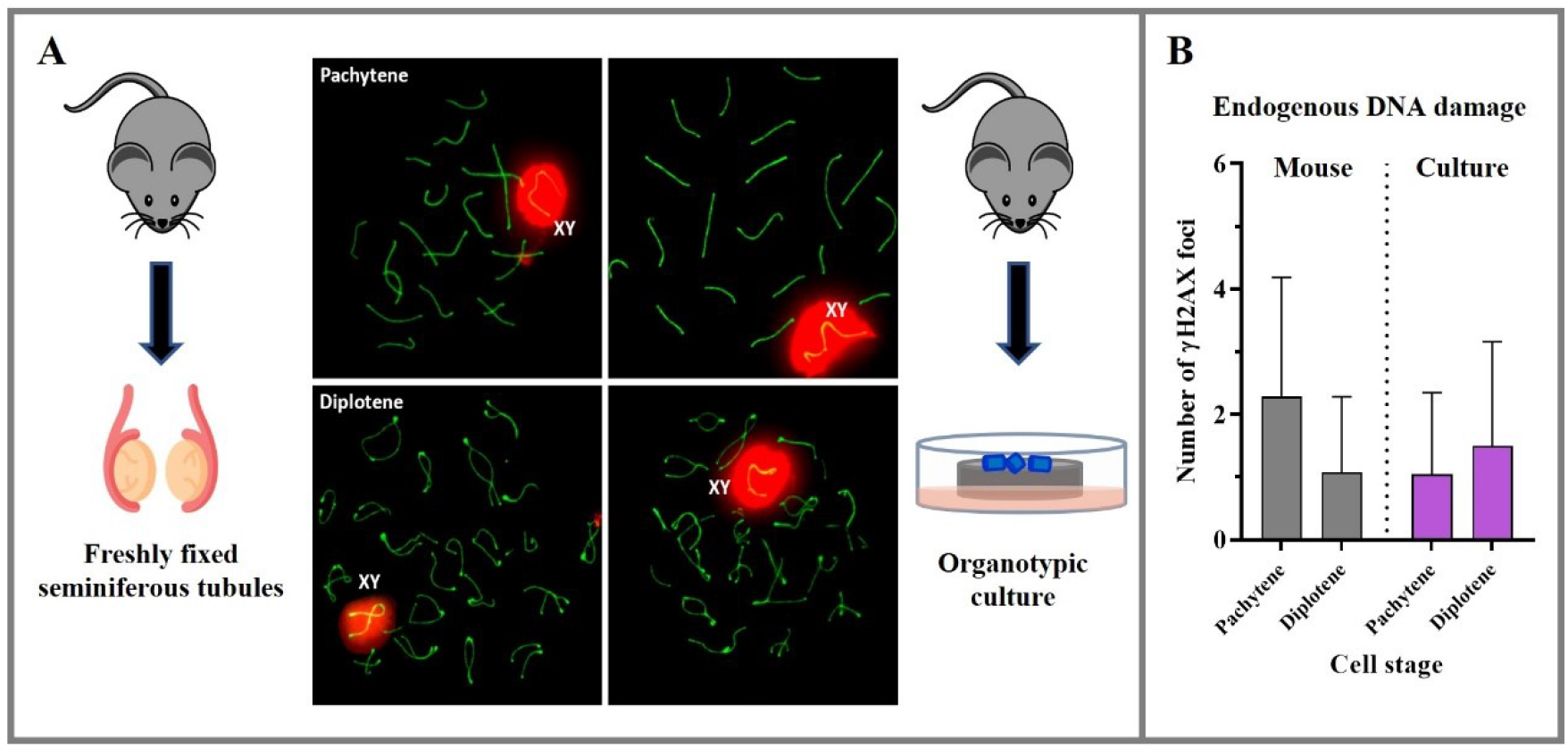
Validation of the pattern of distribution of meiotic markers in organotypic cultures of control seminiferous tubules. (A) Double immunolabelling of SYCP3 (green) and γH2AX (red) of squashed spermatocytes at pachytene and diplotene stages from freshly processed (left) or organotypicly cultured seminiferous tubules (right). (B) Quantitative analysis of the level of endogenous DNA damage detected by γH2AX in all the experimental conditions (n=20 spermatocytes for each condition in three biological replicates).

**Figure 3:**
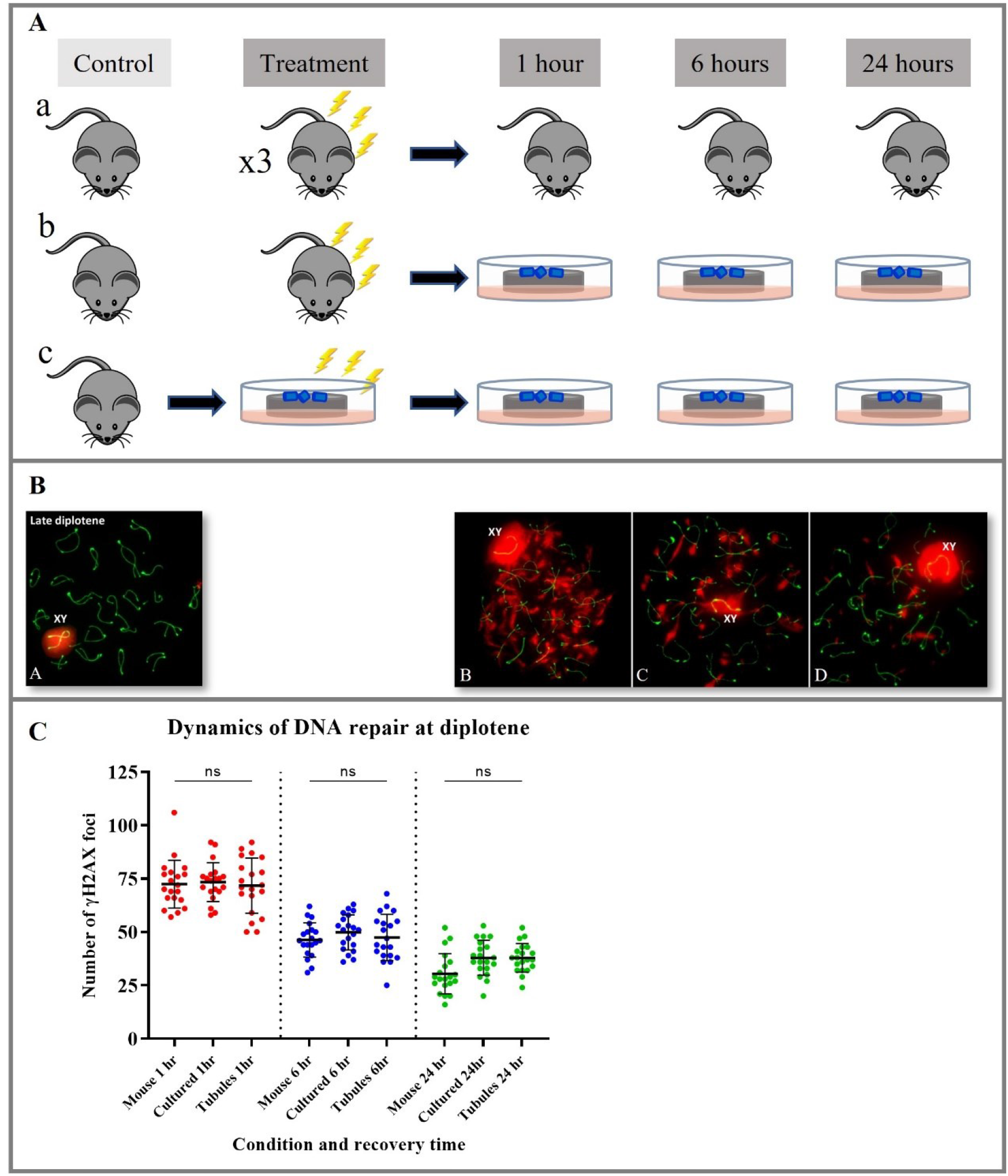
Validation of the DNA damage repair in organotypic cultures of control seminiferous tubules. (A) Experimental approach for the comparison of the DNA damage repair in three experimental conditions (see main text): a) irradiation of alive animals and processing of fresh material (top). This required the use of 4 animals: 1 control, plus 3 irradiated mice processed at different recovery times (1, 6 and 24 hours); b) irradiation of alive animals and subsequent organotypic culture of dissected seminiferous tubules (middle). Seminiferous tubules were processed at the three different recovery times. This approach required the use of two animals: 1 control and one irradiated; and c) irradiation of cultured fragments of seminiferous tubules, which were processed at different recovery times (bottom). This required the use of only one animal. (B) Representative images of the distribution of SYCP3 (green) and γH2AX (red) after immunolabelling on spread spermatocytes in control and different times of recovery post irradiation. The images shown correspond to the irradiation of cultured seminiferous tubules (condition c). (C) Quantitative analysis of the dynamics of DNA damage repair identified by γH2AX foci in the three experimental conditions (a, b and c): mouse (representing approach a), cultured (representing approach b) and tubules (representing approach c) (n=20 spermatocytes for each condition).

## 4. Notes

1. Carry out all procedures at room temperature unless otherwise specified.
2. All instrumentation (petri dishes, well plates, razorblades, tweezes, etc.) must be handled in sterile conditions.
3. When developing the organotypic culture, fragments of the seminiferous tubules should remain over the agarose gel (not immersed in the medium) and they should not move or sink into the bottom of the well.
4. When preparing the material for the squashing technique, the slides covered with poly-L-lysine (Sigma) should be kept covered until use to avoid deposition of dust into the surface.
5. When developing the spreading of spermatocytes technique, fragments of seminiferous tubules extracted for different wells (as long as they were all kept under the same conditions) can be mixed to proceed with cell disaggregation.
6. For functional studies with the use of drugs, it has to be taken into account that the fragments of seminiferous tubules are not immersed in the medium, but rather deposited over the agarose gel absorbing the media from below, therefore requiring higher concentration when developing drug treatments.
7. Although the culture of the seminiferous tubules is a reliable method to carry out *in vitro* meiosis studies, undetermined deviations from normal physiological conditions could arise. In this regard, a recent study suggested that testicular explants could show dysregulation of their transcriptomic profile compared to controls for genes related to inflammation response, insulin-like growth factor and genes involved in steroidogenesis [42]. In contrast, no major differences were found in DNA methylation between mice born after transplantation of long-term cultured spermatogonial stem cells and control, both in F1 and F2 sperm [43]. Therefore, it is advisable to optimize the *in vitro* procedures at the specific conditions for each new study, and each cell type to be examined, before using this approach.

## Acknowledgements

We are very grateful to José A. Suja for his advice and scientific support as some of the cited publications were partly developed and partially funded within his research group [35-38]. We thank all the students that have helped perfectionated the organotypc culture technique presented in this work (Tania Moreno, Irene Mena, Nuria García, Enrique Alfaro, Blanca Tejeda and Inés Hidalgo).

The experiments showed in this work were funded by Ministerio de Ciencia e Innovación (Spain) grants: BFU2009-10987 to J.P. and PID2022-140364NB-I00 to J.P and R.G.; by the Ministerio de Economia y Competitividad (Spain) grant CGL2014-53106-P to J.P., by BIOUAM02-2020 to J.P. and R.G., and by BIOUAM05-2022 to J.P. and R.G.

## Notes

### Competing Interest Statement

The authors have declared no competing interest.

### Summary of Updates

Updated references and incorporation of a new author who9 has collaborated in the manuscript since the first submission

